# ZIKV Induction of Tristetraprolin in Endothelial and Sertoli Cells Post-Transcriptionally Inhibits IFNβ/λ Expression and Promotes ZIKV Persistence

**DOI:** 10.1101/2023.05.03.539309

**Authors:** William R. Schutt, Jonas N. Conde, Megan C. Mladinich, Grace E. Himler, Erich R. Mackow

## Abstract

Zika virus (ZIKV) is a mosquito-borne *Flavivirus* that persistently infects patients, enters protected brain, placental, and testicular compartments, is sexually transmitted, and causes fetal microcephaly *in utero*. ZIKV persistently infects brain microvascular endothelial cells (hBMECs) that form the blood-brain-barrier and Sertoli cells that form testicular barriers, establishing reservoirs that enable viral dissemination. ZIKV persistence requires inhibiting interferon (IFN) responses that direct viral clearance. We found that ZIKV induces IFN-β and IFN-λ in hBMECs but post-transcriptionally inhibits IFN-β/λ expression. IFNβ/λ mRNAs contain AU-rich elements (AREs) in their 3’ untranslated regions which regulate protein expression through interactions with ARE binding proteins (ARE-BPs). We found that ZIKV infection of primary hBMECs induces the expression of the ARE-BP tristetraprolin (TTP) and that TTP is a novel regulator of endothelial IFN secretion. In hBMECs, TTP knockout (KO) increased IFN-β/λ_1_ mRNA abundance and IFN-β/λ_1_ secretion in response to ZIKV infection and inhibited viral persistence. In contrast, TTP expression dramatically reduced IFN-β/λ_1_ secretion in hBMECs. IFN-β/λ_1_ mRNA stability was not significantly altered by TTP and is consistent with TTP inhibition of IFN-β/λ_1_ translation. TTP is similarly induced by ZIKV infection of Sertoli cells, and like hBMECs, TTP expression or KO inhibited or enhanced IFN-β/λ mRNA levels, respectively. These findings reveal a mechanism for ZIKV induced TTP to promote viral persistence in hBMECs and Sertoli cells by post-transcriptionally regulating IFN-β/λ secretion. Our results demonstrate a novel role for virally induced TTP in regulating IFN secretion in barrier cells that normally restrict viral persistence and spread to protected compartments.

**Importance:** Our findings define a novel role for ZIKV induced TTP expression in regulating IFN-β/λ production in primary hBMECs and Sertoli cells. These cells comprise key physiological barriers subverted by ZIKV to access brain and testicular compartments and serve as reservoirs for persistent replication and dissemination. We demonstrate for the first time that the ARE binding protein TTP is virally induced and post-transcriptionally regulates IFN-β/λ secretion. In ZIKV infected hBMEC and Sertoli cells, TTP knockout increased IFN-β/λ secretion, while TTP expression blocked IFN-β/λ secretion. The TTP directed blockade of IFN secretion permits ZIKV spread and persistence in hBMECs and Sertoli cells and may similarly augment ZIKV spread across IFN-λ protected placental barriers. Our work highlights the importance of post-transcriptional ZIKV regulation of IFN expression and secretion in cells that regulate viral access to protected compartments and defines a novel mechanism of ZIKV regulated IFN responses which facilitate neurovirulence and sexual transmission.

## Introduction

Zika virus (ZIKV) is a mosquito-borne neurovirulent *Flavivirus* that uniquely spreads sexually, persists for months in humans and causes encephalitis and fetal microcephaly *in utero* (1-5). In adults, ZIKV is found persistently in saliva, urine, cerebrospinal fluid, and semen causing encephalitis, Guillian-Barre syndrome, and permitting sexual transmission (5-7). During pregnancy, ZIKV spreads across the placenta and *in utero* transmission results in damage to fetal neurons, neural progenitors, and astrocytes that limit brain development and can result in characteristic fetal microcephaly (8, 9). Accordingly, ZIKV infects and crosses endothelial cell (EC), placental, and testicular barriers that normally restrict viral access to the central nervous system, fetal tissues, and semen (1, 7, 10-12). The blood-brain-barrier (BBB) is a neurovascular complex formed by brain microvascular endothelial cells (BMECs), pericytes, and astrocytes that separate the brain from circulating blood constituents and pathogens. Our lab established that ZIKV persistently and non-lytically infects hBMECs, releasing progeny virus both apically and basolaterally without permeabilizing model BBBs (13-15). Additional studies have shown that Sertoli cells that form testicular barriers are persistently infected by ZIKV (16, 17). These findings suggest roles for hBMECs and Sertoli cells as ZIKV reservoirs that facilitate neuroinvasion and sexual transmission.

ECs are a unique cell type with distinct receptors, signaling pathways, and interferon (IFN) directed responses that differentiate them from immune and epithelial cells and tailor their response to viral infection (18, 19). The endothelium is a target for many tick- and mosquito-borne flaviviruses that cause vascular or neurotropic diseases, including Dengue virus (DENV), West Nile virus (WNV), and Powassan virus (POWV) (20-22). ECs produce type I IFN-β and type III IFN-λ in response to viral infections and respond to IFN-⍺/β through cognate type I IFN receptors (IFNARs). ECs lack IFN-λ receptors (IFNλRs) and the addition of IFN-λ to cells fails to inhibit ZIKV infection or induce an interferon stimulated genes (ISGs). Prior treatment of cells with IFN-α/β inhibits ZIKV and other flavivirus infections and DENV induced IFN-α/β secretion following infection of human ECs inhibits DENV spread and ultimately leads to viral clearance (13, 22-24). In contrast, ZIKV persistently infects hBMECs without IFN-β secretion, spreading in monolayers for >9 days and following cellular passage (13, 15). Similar ZIKV persistence has been reported in Sertoli cells, an epithelial cell type that comprises the blood-testes barrier and is hypothesized to facilitate viral entry into the testes and contribute to long term viral shedding and sexual transmission (16, 17, 25). How ZIKV persists within brain and testicular barrier cells is not fully understood however, the ability of ZIKV to restrict IFN responses that prevent viral clearance is likely central to ZIKVs novel persistence, spread, and neurovirulence.

ZIKV has multiple mechanisms of inhibiting IFN responses both upstream and downstream of IFN induction and secretion (26). To impair viral RNA recognition, ZIKV inhibits the activation of retinoic acid-inducible gene I (RIG-I) and melanoma differentiation-associated protein 5 (MDA-5) via expression of several nonstructural proteins and genomic sfRNA (27-32). ZIKV also inhibits the induction of antiviral ISGs directed by IFN receptor activation by inhibiting JAK-STAT2 signaling responses (23, 26, 33). Despite suppression of the IFN induction and IFN signaling pathways, IFN-β and IFN-λ are induced by ZIKV infection of a variety of cell types including hBMECs and Sertoli cells (13, 34, 35). Despite IFN induction in ZIKV infected hBMECs, we found that the expression and secretion of IFN-β/λ in cell supernatants was repressed below the limit of detection and failed to inhibit viral infection or spread (13). Secretory pathways were not globally altered by ZIKV as CCL5, a highly induced, pro-survival chemokine was highly secreted into ZIKV infected hBMEC supernatants. Although most studies evaluate IFN transcripts as surrogates for IFN expression, a discrepancy between IFN-β/λ induction and secretion has also been reported following ZIKV infection of dendritic cells, peripheral blood mononuclear cells, fetal neural progenitor cells, and placental macrophages (34, 36-38). Although a mechanism for post-transcriptional regulation of IFN has not been proposed, the absence of IFN secretion was hypothesized to result from inhibited IFN translation (34).

IFN-β/λ belong to a family of tightly regulated inflammatory cytokines, present at low endogenous levels, that undergo post-transcriptional regulation to rapidly repress secretion (39). Transcript regulation is reportedly mediated by the presence of AU-rich elements (ARE) found in 3’ UTRs of cytokine mRNAs. ARE recognition sequences drive interactions with ARE binding proteins (ARE-BPs) that positively or negatively regulate mRNA translation and stability (39-46). How AREs and ARE-BPs influence IFN-β/λ expression during ZIKV infection has not been investigated.

Studies involving the ARE-BP tristetraprolin (TTP) have largely focused on its role in LPS treated myeloid cells, with little understanding of its role in type I/III IFN regulation or following viral infection. In ECs, expression of the TTP has been shown to regulate the expression of inflammatory cytokines (47, 48). TTP post-transcriptionally inhibits the expression of ARE-containing mRNAs through two separate mechanisms that may function independently or simultaneously (49). TTP can promote the degradation of ARE-mRNAs through deadenylation via recruitment of the CCR4-NOT exonuclease complexes or DCP1/2 decapping enzymes and Xrn1 digestion (50-54). TTP is also capable of translational repression via recruitment of inhibitory translational complexes to mRNAs that interfere with cap binding to eIF4E and translation initiation complex assembly (49, 55-58). How AREs and ARE-BPs influence IFN-β/λ expression during ZIKV infection has not been studied.

Here we investigate ZIKV directed post-transcriptional regulation of IFN expression and define roles for ARE binding proteins in ZIKV persistence in hBMECs and Sertoli cells. In contrast to ARE-BPs AUF1, HuR, and KHSRP, we found that TTP is uniquely induced in ZIKV infected hBMECs and that KO or expression of TTP, respectively, increases or blocks IFN-β/λ secretion. Our findings identify TTP as a ZIKV induced protein in hBMECs and Sertoli cells that regulates the expression of IFN-β and IFN-λ by inhibiting their translation. In addition, we found that ZIKV induces TTP in primary human Sertoli cells and that TTP expression or KO regulates IFN-β/λ secretion. These findings reveal a post-transcriptional mechanism of ZIKV directed IFN regulation resulting from TTP induction following infection and permits ZIKV persistence in cells that normally restrict viral entry into protected compartments.

## Results

### ZIKV prevents IFN secretion in order to persist in hBMECs

We previously reported that ZIKV persistently infects hBMECs, transiently inducing IFNβ and IFNβ/λ_1_ without IFNβ/λ secretion (13). Here we extend these findings and define roles for the ARE-BP TTP in ZIKV directed post-transcriptional regulation of IFN-β/λ in hBMECs. We infected primary hBMECs with ZIKV (PRVABC59) and monitored viral spread and the induction and secretion of IFN-β and IFN-λ_1_. ZIKV titers increased 24 – 72 hpi and monolayers remained persistently infected despite passaging ZIKV infected cells 3 and 6 dpi (Fig. 1A). Following ZIKV infection of hBMECs, we found that both IFN-β and IFN-λ_1_ were transcriptionally induced 1-3 dpi (Fig. 1B, C), but we failed to detect IFN-β or IFN-λ secretion into cell supernatants by ELISA (Fig. 1D, E). In contrast, hBMECs transfected with Poly (I:C) for 24 hours directed the secretion of both IFN-β/λ_1_ (Fig. 1D, E). Concurrently treating hBMECs with IFN-⍺ during ZIKV infection reduced the number of infected cells by 80%, while IFN-λ treated hBMECs had no effect on ZIKV infection (Fig. 1F). Consistent with this, ZIKV persistence and spread in hBMECs functionally demonstrates the absence of IFN-β secretion in ZIKV infected hBMEC supernatants. Validating the presence of IFN-α*/*β, but not IFN-λ, receptors on hBMECs, cellular ISGs (ISG15, MXA) were transcriptionally induced by IFN-⍺, but not by IFN-λ addition to hBMECs, demonstrating that hBMECs only express type I IFNARs and selectively respond to IFN-α*/*β (Fig. 1G, H). Our findings indicate that ZIKV infection of hBMECs post-transcriptionally inhibits IFN-β/λ expression to facilitate ZIKV persistence and spread.

**Figure 1.**
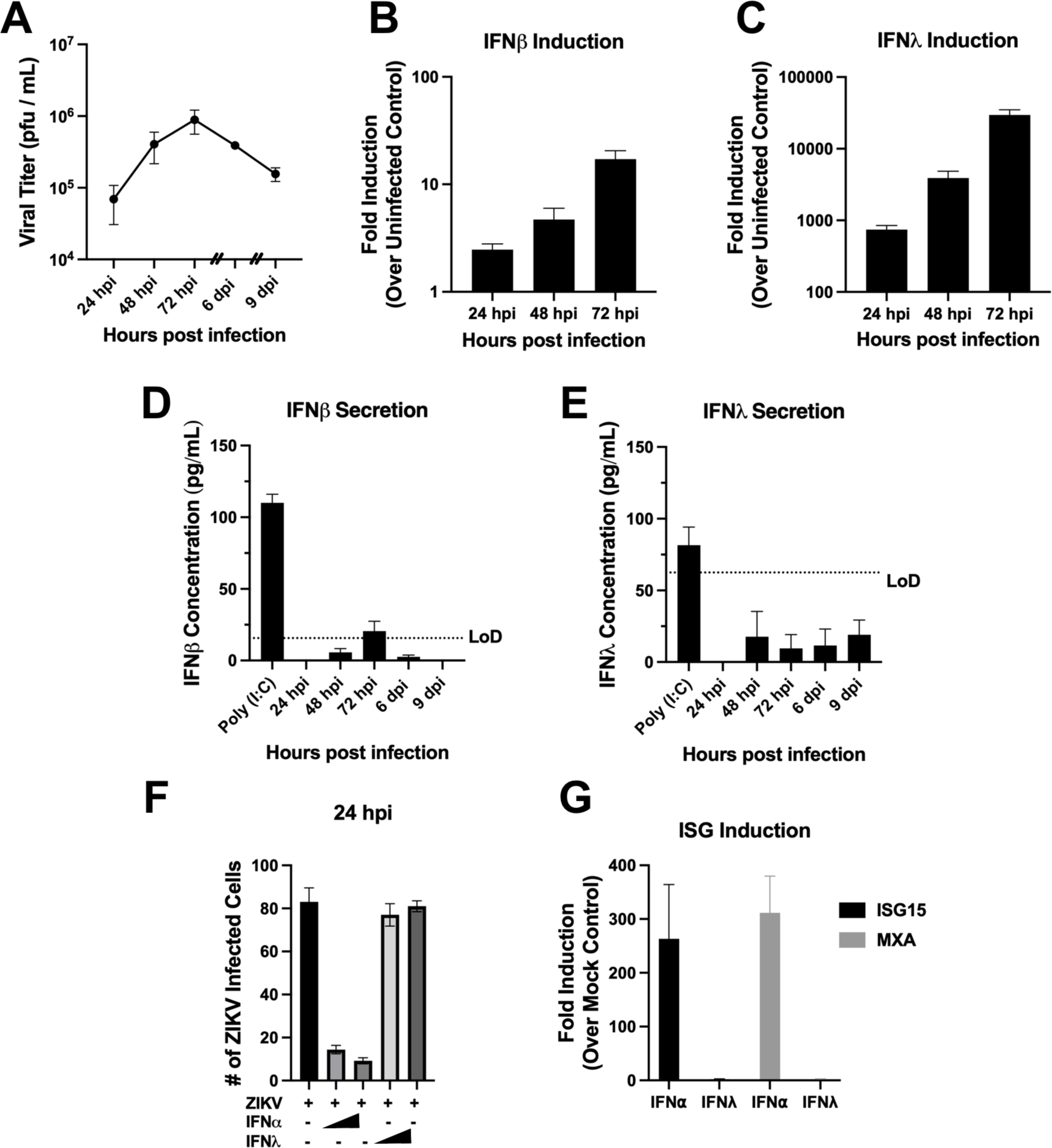
ZIKV prevents IFN secretion in order to persist in hBMECs. (A) Primary hBMECs, were ZIKV infected (MOI = 0.05) and supernatants collected at the indicated times. Dashes on the x-axis indicate when infected hBMECs were trypsinized and passaged (3 and 6 dpi). Viral titer was determined by FFU assay on Vero E6 cells. (B - E) RNA and supernatants from ZIKV infected hBMECs (MOI 0.05) were collected. The induction of IFN-β (B) and IFN-λ_1_ (C) was quantified via qRT-PCR and the concentration of secreted IFN-β (D) and IFN-λ (E) was determined by ELISA. (F, G) WT hBMECs were ZIKV infected (MOI 0.5) with or without simultaneous IFN-⍺ (1000 U/mL or 2000 U/mL) or IFN-λ_1_ (10 ng/mL or 50 ng/mL) addition. The number of ZIKV antigen-positive hBMECs was quantified at 24 hpi using anti-DENV4 HMAF (F) and RNA collected to assess ISG induction (G). Data are represented as the mean ± SEM. Experiments were performed at least 3 times.

### The ARE-BP TTP is induced by ZIKV infection and localized to the cytoplasm

It is widely reported that ZIKV infection transcriptionally induces IFN-β/λ. To explain the discrepancy between IFN mRNA levels and a lack of protein secretion, we hypothesized that ZIKV regulates IFN expression and secretion at a post-transcriptional level. We first performed a miRNA screen of ZIKV infected hBMECs at 24 hpi but failed to detect significant upregulation of any canonical IFN-β/λ targeting miRNAs (data not shown). Based on the potential roles of AREs in cytokine 3’UTRs regulating mRNA expression, we evaluated the expression of ARE-BPs in response to ZIKV infection of hBMECs. IFN-β and IFN-λ contain 3’UTR ARE domains that may regulate IFN expression in a cell type specific manner (39, 40, 46). We initially evaluated expression levels of ARE-BPs HuR, KHSRP, AUF1, and TTP in ZIKV infected hBMECs 1-3 dpi. We found that AUF1, HuR, and KHSRP were expressed constitutively in mock and ZIKV infected hBMECs, with no change in expression following ZIKV infection (Fig. 2A). In contrast, TTP is expressed at extremely low constitutive levels in hBMECs and highly induced by ZIKV 1-3 dpi compared to mock infected cells.

**Figure 2.**
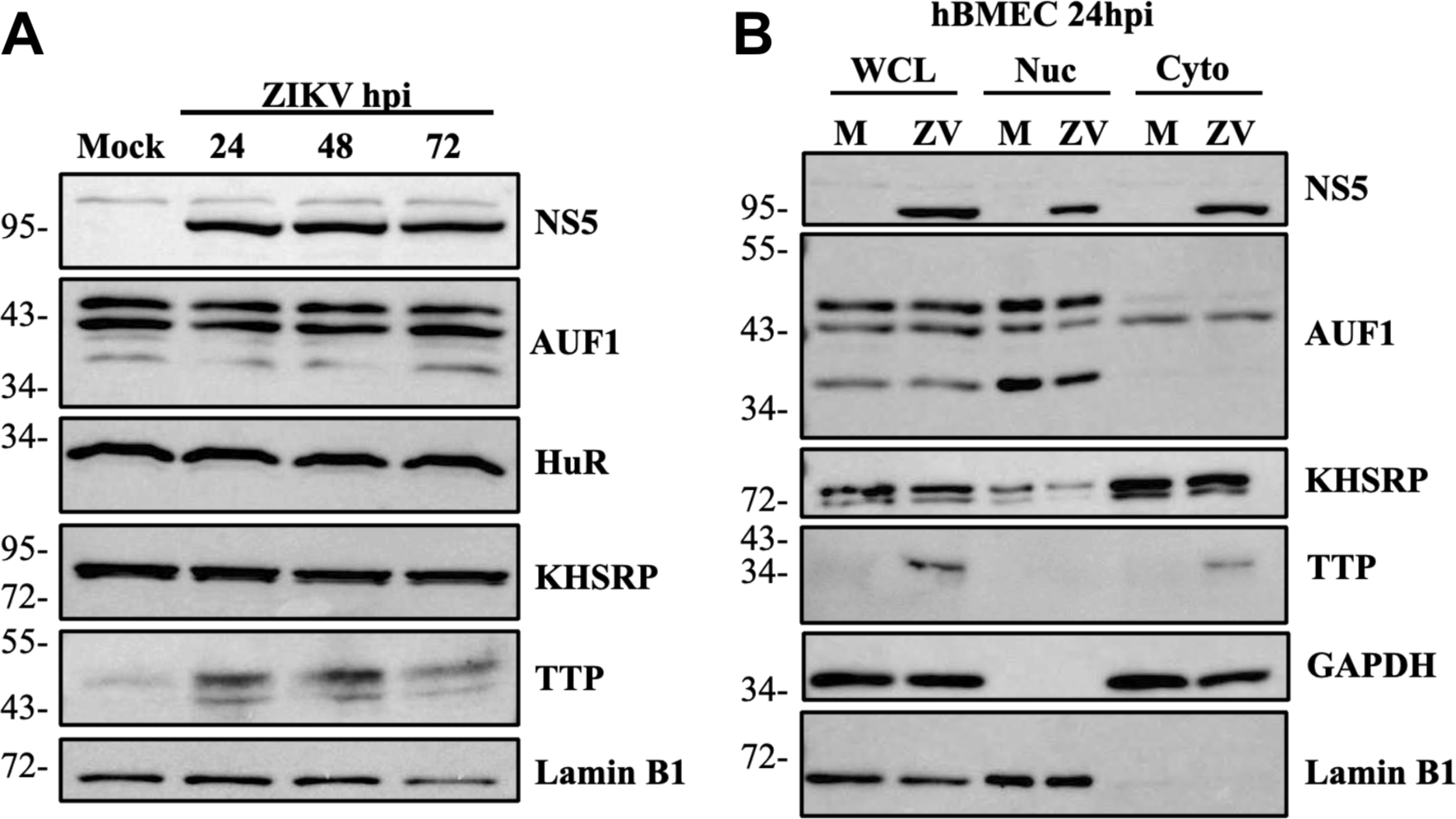
The ARE-BP TTP is induced and localized to the cytoplasm by ZIKV infection. (A) Primary hBMECs were ZIKV infected (MOI 1.0) and lysates collected at the indicated times and the expression of ARE-BPs assessed by Western blot. (B) Mock and ZIKV infected hBMECs (MOI 1.0) were lysed and whole cell lysate (WCL) collected prior to centrifugation and splitting of cytoplasmic (Cyto) and nuclear (Nuc) fractions. Lysates were Western blotted for cytoplasmic GAPDH and nuclear lamin B1 protein markers to confirm equal loading. Western blots are representative of multiple independent experiments.

Regulation of mRNAs by ARE-BPs is linked to protein activation that directs localization from the nucleus to the cytoplasm (59-62). We evaluated ARE-BP localization in ZIKV versus mock infected cells 24 hpi and observed no change in the nuclear or cytoplasmic localization of AUF1 or KHSRP in mock or ZIKV infected hBMECs (Fig. 2B). The absence of TTP in mock infected cells prevents analysis of a change in TTP localization, but the exclusive localization of TTP in the cytoplasm of ZIKV infected hBMECs indicates the unique induction of activated TTP in ZIKV infected hBMECs (Fig 2B).

### ZIKV induction of TTP is independent of IRF3 and IFN

TTP induction has primarily been studied in myeloid cells in response to inflammatory LPS and TNF-⍺ stimuli, while the role of IFNs is unclear (59, 63). As IFN-β/λ secretion is restricted in ZIKV infected hBMECs, we compared TTP induction in response to ZIKV infection or stimulation with IFN-α*/*λ. hBMECs were ZIKV infected, treated with 1000U/mL of IFN-⍺, or 50ng/mL IFN-λ_1_ for 24 hours prior to analysis of TTP induction by qRT-PCR (Fig. 3A). We found that ZIKV virus infection, but not IFN-⍺/λ treatment, induced TTP in hBMECs. These findings are consistent with a previous study indicating that knockdown of RIG-I partially reduced TTP induction in response to RNA transfection (64). To determine if TTP induction is mediated by IRF3 activation following ZIKV infection we transduced hBMECs to express a dominant negative IRF3 (IRF3Δ60) that lacks the ability to induce IFN-β (65). In response to ZIKV infection IFN-β was induced in WT hBMECs but not IRF3Δ60-hBMECs and IRF3Δ60 expression dramatically reduced IRF3 directed ISG expression (Fig. 3B, S1). Protein analysis revealed that TTP expression was induced by ZIKV infection of both WT and IRF3Δ60 hBMECs (Fig. 3C). These findings demonstrate that ZIKV infection of endothelial cells transcriptionally induces TTP expression independently of IRF3 or IFN signaling responses.

**Figure 3.**
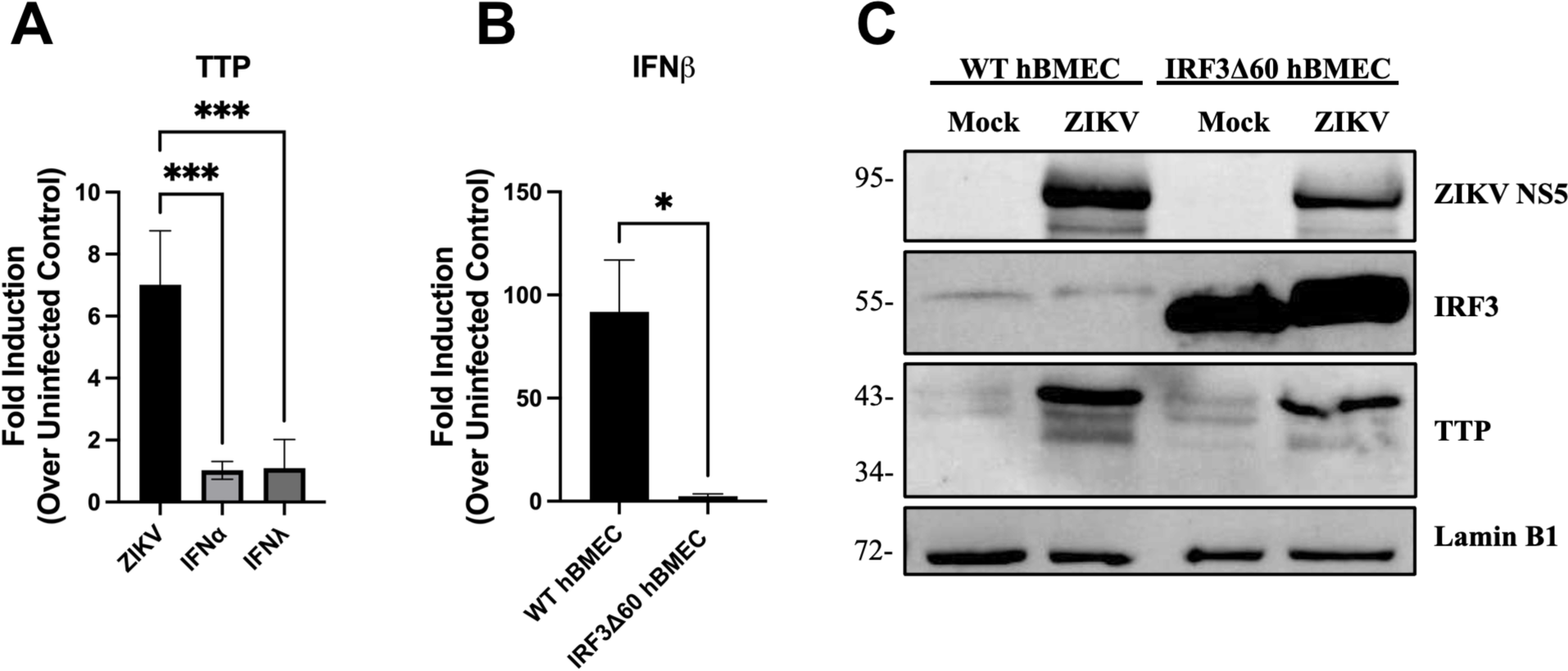
ZIKV induction of TTP is independent of IRF3 and IFN. (A) Primary hBMECs were ZIKV infected (MOI 10), treated with exogenous IFN-⍺ (1000 U/mL), or IFN-λ_1_ (50 ng/mL) for 24 hpi prior to RNA collection and analysis of TTP transcriptional induction by qRT-PCR. Statistical significance was determined by one-way ANOVA (B, C) Primary hBMECs, and hBMECs expressing dominant negative IRF3Δ60 were ZIKV infected (MOI 10) with RNA and protein lysates collected 1-2 dpi. Transcriptional induction of IFN-β was determined by qRT-PCR (B) and significance determined via unpaired student’s T test. The expression of IRF3 and TTP analyzed by Western blot (C). Data are represented as the mean ± SEM, asterisks indicate statistical significance (*, P ≤ 0.05; ***, P ≤ 0.001). Experiments were performed 3 times.

### TTP expression inhibits IFN-β/λ expression

While TTP has been shown to inhibit the expression of some ARE-containing cytokines, regulation of IFNs remains understudied and has only been addressed in immune cells. To evaluate the effect of TTP on IFN-β/λ_1_ induction and secretion in hBMECs, we generated TTP CRISPR-Cas9 KO and doxycycline-induced TTP overexpressing hBMECs (Fig. S2). Following Poly (I:C) transfection, TTP KO hBMECs exhibited higher induction of IFN-β/λ_1_ that was dramatically reduced by TTP overexpression (Fig 4A, B). The enhanced IFN mRNA abundance corresponded with a significant increase in IFN-β/λ_1_ secretion by TTP KO cells 24 hours post transfection (Fig 4C, D). These findings demonstrate direct and novel IFN-β/λ regulation by TTP in endothelial cells and suggest a mechanism for virally induced TTP to modulate inflammatory responses of the endothelium.

**Figure 4.**
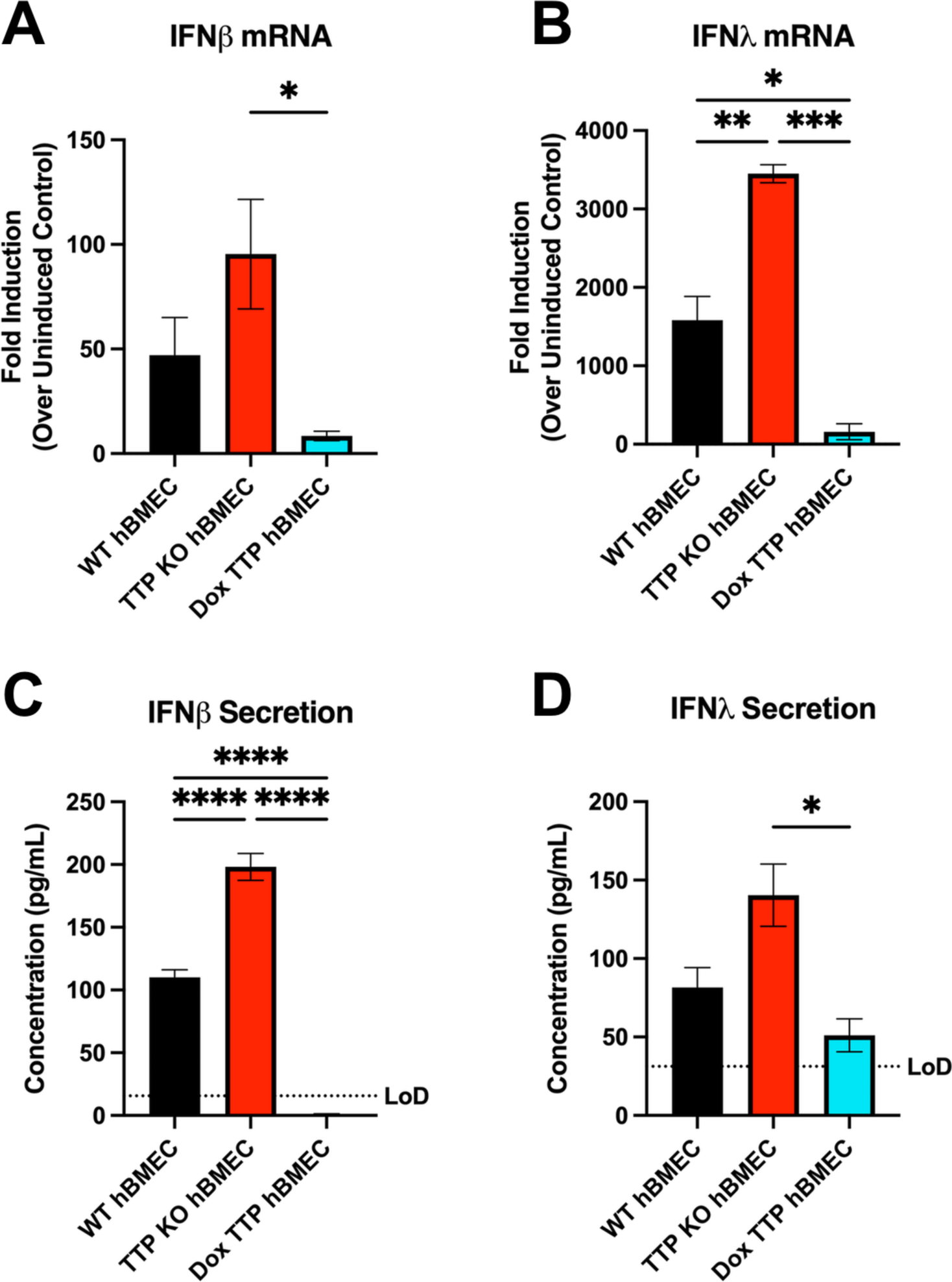
TTP expression inhibits IFN-β/λ expression. Primary hBMECs, TTP-KO hBMECs, and TTP expressing hBMECs were ZIKV infected (MOI 10) for the indicated times. TTP expression was doxycycline induced (200 ng/mL) throughout the course of the experiment. Transcriptional induction of IFN-β (A) and IFN-λ_1_ (B) was determined via qRT-PCR. Infected hBMEC supernatants were measured by ELISA for IFN-β (C) and IFN-λ (D). Data are represented as the mean ± SEM, asterisks indicate statistical significance as determined by one-way ANOVA (*, P ≤ 0.05; ***, P ≤ 0.001, ****, P ≤ 0.0001). Experiments were performed at least 3 times.

### ZIKV persistence is suppressed in TTP KOs and enhanced in TTP expressing hBMECs

We determined if TTP KO or TTP expression in hBMECs affected ZIKV replication and persistence in hBMECs. In a synchronous ZIKV infection of WT, TTP KO, and TTP expressing hBMECs, we found that titers rapidly reached maximal levels (1 x 10^6^/ml) 1-2 dpi with little difference between cell types (Fig. 5A). However, this single step kinetic experiment only evaluates initial viral output that does not require ZIKV to spread to secondary cells. To assess roles for TTP in viral persistence, we passaged ZIKV infected WT, TTP KO, and TTP expressing cells and monitored viral titers 6 to 9 dpi. We observed a significant reduction in persistently infected ZIKV titers in TTP KO hBMECs at 6 and 9 dpi, versus hBMECs that express TTP (WT or TTP expressing). In support of TTP regulated ZIKV persistence, staining of ZIKV infected hBMECs 6-9 dpi revealed reduced ZIKV spread in TTP KO cells versus WT or TTP expressing hBMECs (Fig. 5B). Collectively, these finding suggest that ZIKV induced TTP expression suppresses IFN secretion and facilitates persistent viral spread.

**Figure 5.**
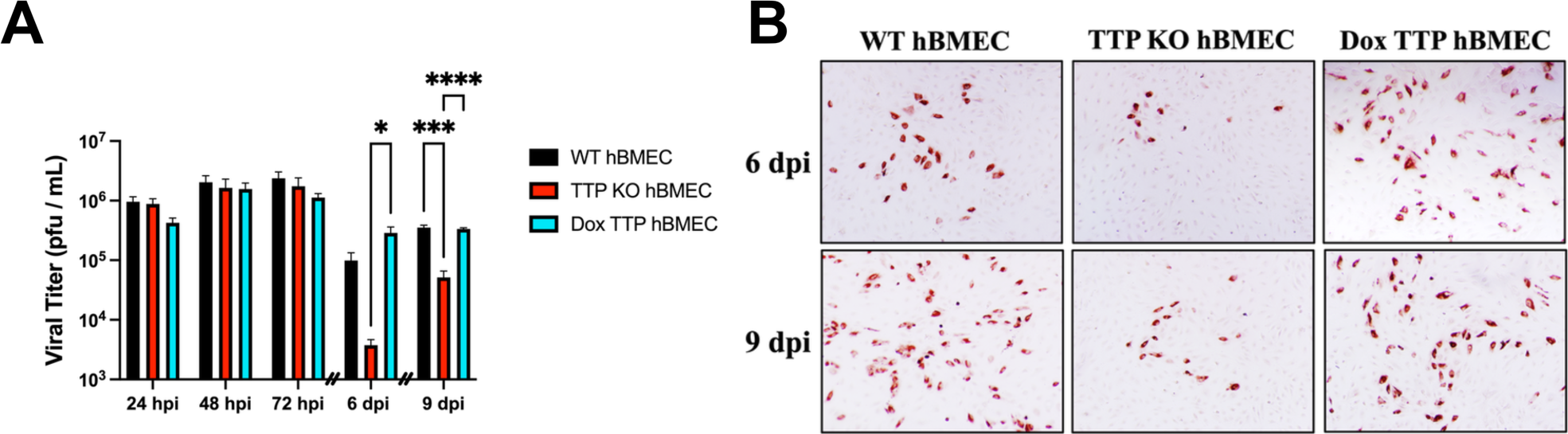
ZIKV persistence is suppressed in TTP KOs and enhanced in TTP expressing hBMECs. (A) Primary hBMECs, TTP-KO hBMECs, and TTP expressing hBMECs were ZIKV infected (MOI 1) and supernatants collected and titered on Vero E6 cells at the indicated times. Dashes on the x-axis indicate when infected hBMECs were trypsinized and passaged (3 and 6 dpi). Data are represented as the mean ± SEM, asterisks indicate statistical significance as determined by two-way ANOVA (*, P ≤ 0.05; ***, P ≤ 0.001, ****, P ≤ 0.0001). Experiments were performed 4 times. (B) Infected hBMECs were fixed and ZIKV positive cells stained with anti-DENV4 HMAF at 6 and 9 dpi following one and two cellular passages respectively.

In order to assess IFN-β/λ_1_ induction and secretion during ZIKV infection we infected hBMECs (MOI 10) and evaluated IFN-β/λ responses in WT, TTP KO, and TTP expressing hBMECs. In TTP KO cells, IFN-β/λ_1_ was induced 2-4 fold over WT hBMECs 1-3 dpi. In contrast, TTP expressing cells dramatically repressed IFN-β/λ_1_ induction 5-10 fold versus WT or TTP KOs (Fig. 6A, B). The secretion of IFN-β was notably enhanced in ZIKV infected TTP KO hBMECs 1-3 dpi, with TTP expressing hBMECs repressing IFN-β secretion to levels 5-30 fold less than WT or TTP KO cells (Fig. 6C – E). A similar trend of IFN-λ_1_ secretion 2-3 dpi in TTP KOs and reduced IFN-λ_1_ secretion in TTP expressing cells was observed but diminished. These findings reveal a novel mechanism of ZIKV directed post-transcriptional IFN regulation directed by the induction of the ARE-BP TTP and demonstrates a novel role for ZIKV induced TTP in regulating viral persistence and spread in endothelial cells.

**Figure 6.**
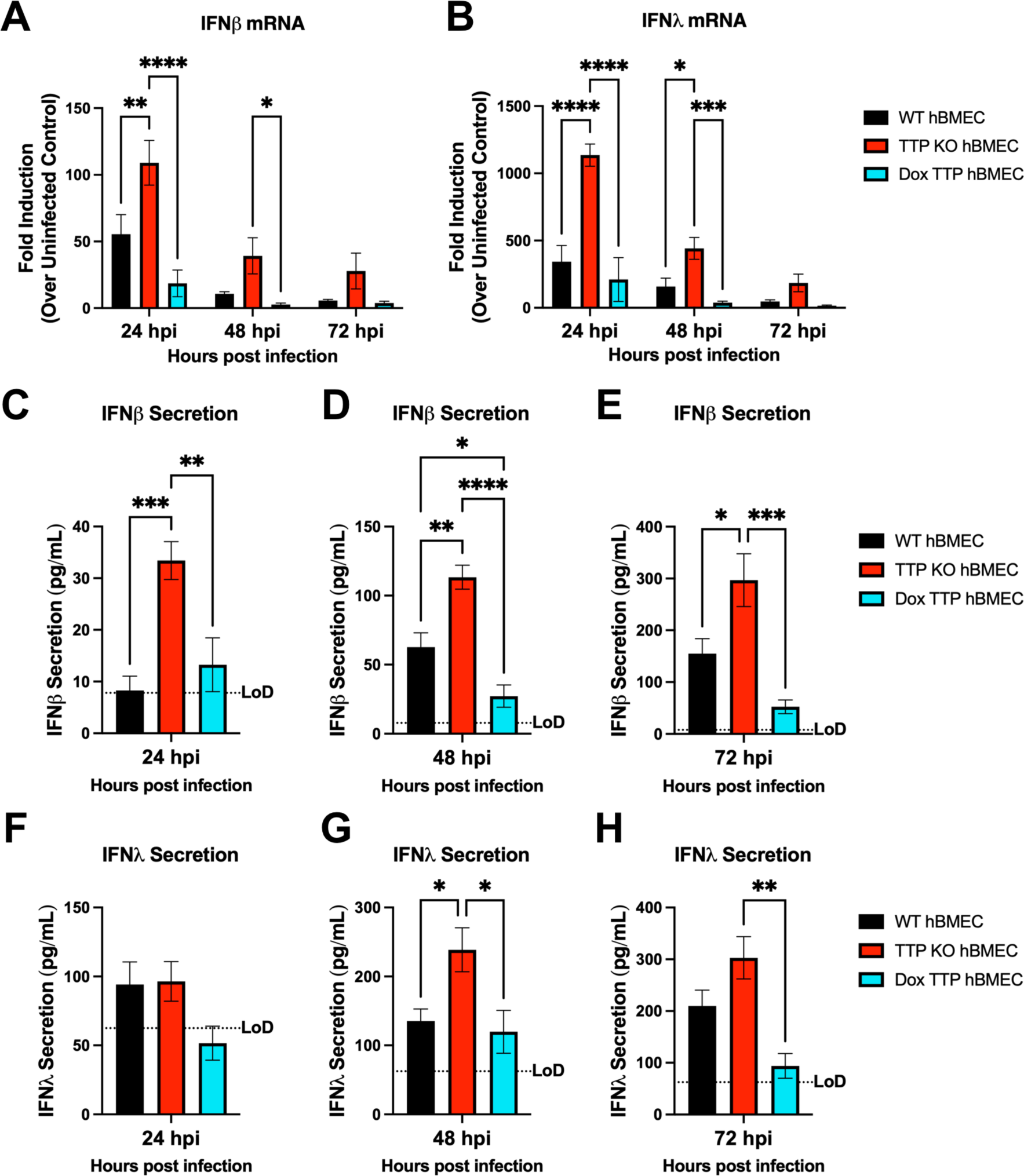
IFN induction and secretion is enhanced during ZIKV infection of TTP KO hBMECs. Primary hBMECs, TTP-KO hBMECs, and TTP expressing hBMECs were ZIKV infected (MOI 10) and cellular RNA and supernatants harvested at the indicated times. IFN-β (A) and IFN-λ_1_ (B) mRNA levels were determined by qRT-PCR. The concentration of IFN-β (C – E) and IFN-λ (F – H) was determined from infected supernatants by ELISA (R&D systems). Data are represented as the mean ± SEM, asterisks indicate statistical significance as determined by two-way ANOVA (*, P ≤ 0.05; **, P ≤ 0.002; ***, P ≤ 0.0002, ****, P ≤ 0.0001). Experiments were performed 4 times.

### TTP KO does not alter IFN mRNA degradation

In immune cells, TTP has been shown to post-transcriptionally repress cytokine expression through TTP complexes that both translationally repress protein expression from ARE containing mRNAs or by reducing the stability of ARE-mRNA transcripts. To differentiate TTP mechanisms of action, we evaluated the rate of IFN-β/λ mRNA degradation in WT and TTP KO cells and compared the effect of TTP to the canonical TTP destabilizing target IL-6. WT hBMECs and TTP KO hBMECs were ZIKV infected (MOI 10) and were treated with actinomycin D (ActD) at 20 hpi to block new RNA transcription and assayed for changes in IFN-β/λ and IL6 mRNA abundance (2-4 hours post ActD addition) by qRT-PCR. In TTP KO hBMECs, there was a noted reduction in the rate of IL6 mRNA degradation that reflects the role of TTP in reducing IL6 mRNA stability in immune cells (Fig. 7C) (66). In contrast, in ZIKV infected hBMECs TTP KO did not significantly alter the rate of IFN-β or IFN-λ_1_ mRNA degradation compared to WT hBMECs (Fig. 7A, B). These findings are consistent with TTP inhibiting IFN-β/λ secretion through the post-transcriptional inhibition of IFN-β/λ_1_ mRNA translation in hBMECs.

**Figure 7.**
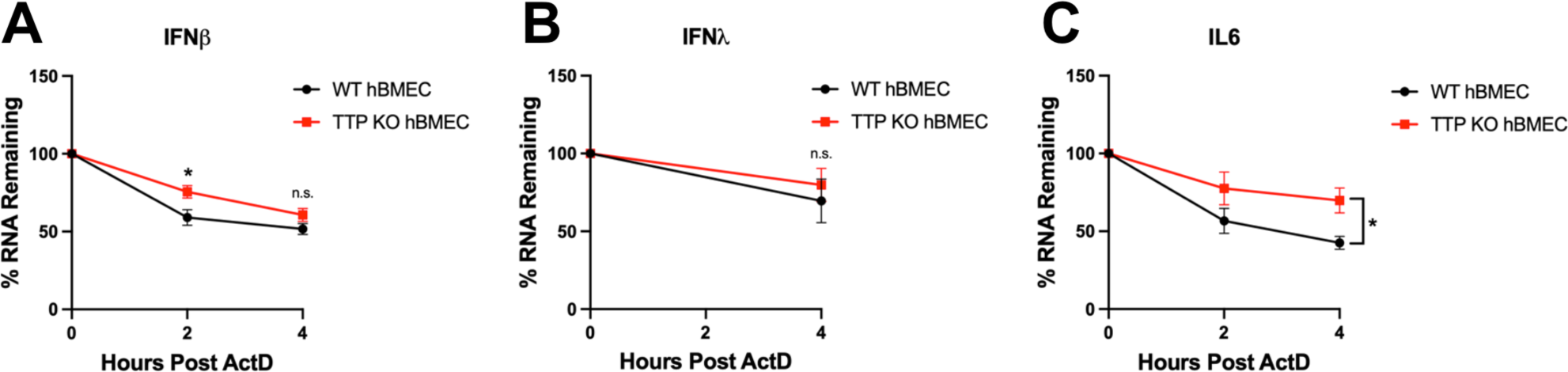
TTP KO does not alter IFN mRNA degradation. Primary hBMECs and TTP-KO hBMECs were ZIKV infected (MOI 10) for 20 hours prior to the addition of actinomycin D (5µg/mL). RNA was collected 0, 2, and 4 hours post ActD addition and the levels of IFN-β (A), IFN-λ_1_ (B), and IL-6 (C) mRNAs analyzed via qRT-PCR and normalized to β-actin. Data are represented as the mean ± SEM, asterisks indicate statistical significance as determined by two-way ANOVA (*, P ≤ 0.05). Experiments were performed 3 times.

### TTP regulates IFN-β/λ expression in human Sertoli cells

ZIKV persistently infects human Sertoli cells (hSerC) that protect testicular compartments, and viral persistence in this cellular niche is likely to enhance ZIKV sexual transmission (16, 17, 67). Sertoli cells produce both IFN-β and IFN-λ_1_ and are sensitive to exogenous type I IFN treatment (Fig. 8A). Like hBMECs, the addition of IFN-λ_1_ to Sertoli cells does not restrict ZIKV replication or induce ISGs (Fig. 8A–C, Fig. 1G, H). We initially determined if ZIKV infection directs ARE-BP responses that inhibit IFN-β/λ_1_ secretion in hSerCs. Primary hSerCs were synchronously infected with ZIKV and the expression of ARE-BPs was monitored 1-2 dpi. Like hBMECs, hSerCs showed no alteration in HuR, KHSRP, or AUF1 expression following ZIKV infection (Fig. 8D). In contrast, TTP expression was dramatically increased in ZIKV infected Sertoli cells 1-2 dpi (Fig 8D). To evaluate the role of TTP in regulating IFN-β/λ_1_ mRNA levels in Sertoli cells, we generated TTP-KO and TTP expressing hSerCs. ZIKV infected TTP KO hSerCs demonstrated a dramatic increase in IFN-β/λ_1_ mRNA levels compared to ZIKV infected WT controls (Fig. 8E, F). Conversely, ZIKV infection of TTP expressing hSerCs reduced IFN-β/λ_1_ mRNA levels by 60-85% when compared to WT or TTP-KO hSerCs (Fig. 8E, F). These findings demonstrate that ZIKV induced TTP regulates IFN-β/λ expression in Sertoli cells, key testicular barrier cells that may serve as reservoirs for ZIKV dissemination contribute to the unique ability of ZIKV to be sexually transmitted. ZIKV directed TTP induction and IFN regulation reveals a fundamental ZIKV persistence mechanism that prevents viral clearance from crucial physiological barriers.

**Figure 8.**
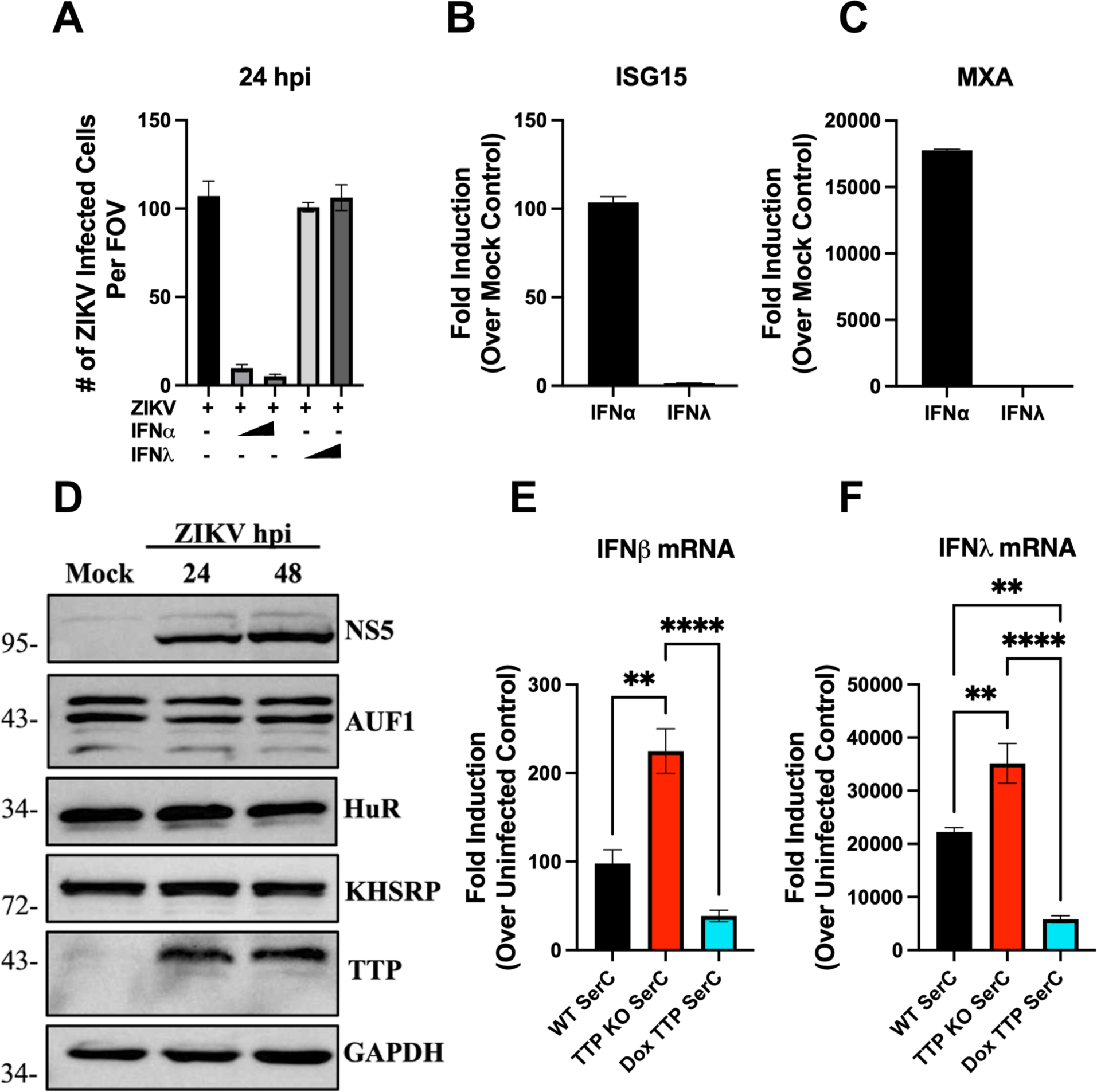
TTP regulates IFN-β/λ expression in human Sertoli cells. (A - C) Primary hSerCs were ZIKV infected (MOI 0.5) with or without simultaneous IFN-⍺ (1000 U/mL or 2000 U/mL) or IFN-λ_1_ (10 ng/mL or 50 ng/mL) addition. ZIKV antigen-positive hSerCs were quantified 24 hpi using anti-DENV4 HMAF (A) and RNA collected to assess ISG induction (B, C). (D) Primary hSerCs were ZIKV infected (MOI 10) and lysates collected at indicated times for analysis of ARE-BP expression by Western blot. (E – F) Western blot is representative of 2 experiments. ZIKV infected (MOI 10) primary hSerCs, TTP KO-hSerCs, and TTP expressing hSerCs were assayed for RNA levels 24 hpi by qRT-PCR analysis of IFN-β (E) and IFN-λ (F) mRNAs. Data are represented as the mean ± SEM, asterisks indicate statistical significance as determined by one-way ANOVA (*, P ≤ 0.05; **, P ≤ 0.002; ***, P ≤ 0.0002, ****, P ≤ 0.0001). Experiments were performed at least 3 times unless otherwise noted.

## Discussion

ZIKV is distinguished from other flaviviruses by its ability to persist in patients for up to 6 months, cause fetal microcephaly *in utero*, and be sexually transmitted (1-4, 68). To accomplish this, ZIKV persistently infects endothelial, trophoblast, and Sertoli cells that serve as barriers to immune protected brain, testicular, and placental compartments (10, 69-72). ZIKV persistence provides a reservoir for ZIKV dissemination and spread across tissue restrictive barriers (13). We previously reported that ZIKV persistently infects blood-brain-barrier endothelial cells, spreading basolaterally from polarized hBMECs, a mechanism consistent with CNS entry. ZIKV persistence in hBMECs requires the autocrine activation of pro-survival CCL5 responses and the inhibition of IFNβ secretion (15). hBMECs secrete Type I (IFN-β) and Type III (IFN-λ) IFNs, however, as hBMECs lack IFNλRs and fail to respond to IFN-λ, ZIKV infection of hBMECs is only restricted by IFN-⍺/β pre-treatment. Despite transient induction of IFN-β/λ in infected hBMECs, ZIKV infected hBMECs failed to secrete detectable or functional IFN-β/λ and lead us to hypothesize ZIKV post-transcriptionally inhibits IFN-β/λ expression (13).

ZIKV impairs IFN induction and IFN receptor responses through multiple mechanisms. ZIKV NS3/NS5, additional NS proteins, and sfRNA reportedly suppress RIG-I/MDA-5 signaling pathway responses that induce IFN mRNA transcription (26-31). ZIKV also targets JAK/STAT signaling pathways downstream of IFNAR to restrict the induction of antiviral ISGs. NS2B3 degrades Jak kinase, preventing downstream STAT phosphorylation, while ZIKV NS5 inhibits STAT phosphorylation and promotes human STAT2 degradation (23, 26, 33).

Despite ZIKV regulation of IFN induction, several reports note that IFN induction is dissociated from IFN secretion (34, 36-38). ZIKV infected PBMCs induced IFN transcripts but failed to secrete IFN-⍺ and IFN-λ, and ZIKV infected placental macrophages, fetal neural progenitors, and dendritic cells, similarly fail to secrete IFNs (34, 36-38). In macrophages a comparison of IFN cDNA synthesis (hexamer versus oligo-dT) suggested IFN mRNA stability was unchanged by ZIKV infection, and from this translational IFN regulation was inferred (34). Collectively, IFN induction without IFN secretion following ZIKV infection was noted in immune cells, where TTP expression in response to LPS is well characterized, and mirror our own observations of ZIKV inhibiting IFN secretion in hBMECs at a translational level (34, 36-38).

ZIKV infection reportedly induces the expression of miRNAs that have been shown to regulate the production of ISGs, cell death, and chemotaxis and represent a potential mechanism of post-transcriptional IFN regulation (73-75). Although ZIKV induction of miRNAs targeting IFN-β/λ have not been reported, the distantly related *flavivirus* hepatitis C virus (HCV) induces miRNAs targeting IFN-λ to restrain IFN signaling and promote viral persistence in hepatocytes (76, 77). We initially assessed miRNAs induced in ZIKV infected hBMECs at 24 hpi but found no significant upregulation of canonical IFN targeting miRNAs (mir-34a, mir-145, or let-7b), with the IFN-β targeting miRNA, miR-34a, induced by only 1.61 fold (p-value = 0.06) (78).

In HCV pathogenesis, IFN-λ ARE-mediated inhibition has been implicated. In the 3’ UTR of *IFNL3,* an ARE polymorphism was identified that interfered with ARE sequence recognition, ultimately enhancing IFN-λ_3_ production and conferring cellular resistance to HCV (77, 79, 80). Both *IFNB1* and *IFNL1* transcripts produced by ZIKV infected hBMECs contain AREs in their 3’ UTRs (46, 81). This led us to hypothesize the AREs in IFN-β/λ mRNAs as targets of ZIKV post-transcriptional regulation and potential roles for ARE-BPs in ZIKV persistence in hBMECs. During ZIKV infection, ARE-BPs have primarily been observed as markers of stress granule localization but ARE-BPs as determinants of spread and persistence have remained understudied (82-84). Knockdown of HuR, an ARE-BP that stabilizes IFN mRNAs, was shown to increase ZIKV titers, although the effect on IFN expression was not assessed (40, 84).

Due to reports of ARE-BPs regulating IFN mRNAs, we assessed the expression and localization of TTP, KHSRP, AUF1, and HuR in ZIKV infected hBMECs (39). We found that only TTP was induced by ZIKV infection and that TTP was localized to the cytoplasm. Importantly, ZIKV directed TTP expression was independent of IRF3 directed IFN induction and TTP was not induced in response to IFN addition to cells. How ZIKV triggers TTP induction independent of IRF3 is unclear and warrants further study, but likely involves regulation by p38 MAPK. ZIKV infection reportedly activates p38 MAPK, a key regulator of LPS directed TTP induction, activity, and protein stability (50, 59, 85, 86).

IFN regulation remains an understudied aspect of TTP biology and has not been addressed in the context of viral infections. Impaired IAV replication following TTP knockdown was attributed to an increase in COX-2 protein, however the effect of TTP on IFN production was not assessed (64). Two studies have implicated IFN-β and IFN-γ as targets of TTP repression in LPS treated murine macrophages and T cells respectively, however TTP regulation of IFN has not been assessed in virally infected, non-immune human cells (87, 88). We observed that TTP expression in hBMECs and Sertoli cells regulates IFN-β/λ mRNA abundance and protein expression in response ZIKV infection. These cells comprise critical physiological barriers protecting brain and testicular tissues respectively and the ability to limit IFN responses likely fosters viral persistence and spread into these immune privileged compartments. Apart from this study, the role of TTP in regulating IFN responses that restrict viral infection and spread in human cells has yet to be studied. In addition to TTP expression in brain, testicular, placental, and ovarian tissues, TTP is the most highly expressed in cervical tissue (89). As IFN-λ secretion protects the female reproductive track and plays a critical role in preventing viral spread across placental barriers, TTP expression may suppress IFN production during ZIKV infection to permit viral spread and congenital fetal infection (90, 91). In these barrier settings, TTP regulated IFN-β/λ responses are likely to be critical to ZIKV sexual transmission, persistence, and *in utero* spread to fetal tissues.

To determine if IFN-β/λ mRNA and secreted protein levels are altered due to TTP destabilization of IFN mRNAs, we assessed IFN-β, IFN-λ_1_, and IL-6 degradation rates following addition of actinomycin D. The ability of TTP to promote IL-6 mRNA degradation is well characterized in immune cells in response to LPS (66, 92). However, TTP regulation of IFNs is less clear with TTP expression enhancing IFN-β mRNA degradation 1 hour, but not 4 hours, post LPS treatment, in murine BMDMs (88). In contrast, KO of a TTP activator, DUSP1, stabilized IFN-β mRNA only at 4 hours post LPS treatment, suggesting that TTP activity is temporally regulated (88). We assessed IFN mRNA stability at 24 hpi because it coincides with the peak of transient IFN induction and TTP expression following a synchronous ZIKV infection of hBMECs. In TTP KO hBMECs, we observed no significant alteration to the IFN-β/λ_1_ mRNA stability but did detect faster degradation of IL-6 mRNA. While we cannot discount altered IFN mRNA stability at other times post ZIKV infection, our findings are consistent with ZIKV induced TTP repressing the translation of IFN-β/λ mRNAs and ultimately suppressing IFN secretion.

TTP has multiple mechanisms of repressing ARE mRNA translation and has been shown to directly form complexes with poly(A)-binding protein and the inhibitory eukaryotic initiation factor 4E2 (57, 58, 93). Recruitment of eIF4E2 to ARE mRNAs prevented their association with eIF4E, a scaffold protein required for assembly of the translation initiation complex leading to the inhibition of cap-dependent translation (57, 58, 93). TTP also reportedly interacts with GYF2 to recruit the cap binding translational repression complex 4EHP that competes for cap binding with eIF4E (56). However, the formation of translation regulating TTP complexes with IFN mRNAs and has not been addressed in endothelial cells, Sertoli cells, or trophoblasts that normally protect tissues from viral access (57, 58, 93).

Overall, our findings characterize a novel role for TTP in suppressing IFN-β/λ production in primary human BMECs and Sertoli cells in response to ZIKV infection. We found that TTP is induced by ZIKV independent of IRF3 and IFN activity, and that TTP post-transcriptionally inhibits IFN-β/λ protein expression and secretion despite transcriptional induction. These novel findings demonstrate that TTP regulation of IFN responses contributes to ZIKV spread and persistence and provides an additional mechanism for induced TTP to regulate IFN-λ expression, a critical determinant of ZIKV placental transmission (94). Our findings underscore the complexity of the IFN response to ZIKV and highlight a new post-transcriptional mechanism of innate immune interference that prevents IFN secretion and fosters ZIKV persistence and spread across key tissue barriers.

## Acknowledgements

We thank Patrick Hearing, Nancy Reich-Marshall, Laurie Krug, Daniel Salamango, Elena Gorbunova, and William A. Schutt, Jr. for their helpful discussions and manuscript feedback. This work was supported by funding from a DOD TBDRP Idea Development Award W81XWH2210702, National Institutes of Health grants: NIAID R01AI12901005, R21AI13173902, R21AI15237201, RO1AI027044, T32AI007539, and a Stony Brook University Seed Grant. The funders had no role in study design, data collection and interpretation or the decision to submit the work for publication. We declare no conflict of interest.

## Methods and Materials

### Cells

C6/36 cells (ATCC CRL-1660) were grown in Dulbecco’s modified Eagle’s medium (DMEM) supplemented with 10% FBS, penicillin (100 µg/ml), streptomycin sulfate (100 µg/ml), and amphotericin B (50 µg/ml; Mediatech) at 28°C with 5% CO_2_. Vero E6 (ATCC CRL 1586) and HEK293T (ATCC) were grown in DMEM as above with 8% FBS. Human brain microvascular endothelial cells (hBMEC) were purchased from Cell Biologics (H-6023) and grown in Endothelial Cell Growth Basal Medium-2 (EBM-2) with SingleQuots (Lonza) at 37°C in 5% CO2. hBMECs were discarded upon reaching passage 12. Human Sertoli cells were purchased from ScienCell (#4520) and grown as above in Sertoli cell media (#4521).

### Virus

ZIKV (PRVABC59) was obtained from the ATCC (VR-1843) and passaged in C6/36 cells according to manufacturer’s instructions. Infectious ZIKV stocks (5 x10^5^ FFU/mL) were generated by inoculating confluent T75 flasks of C6/36 cells for 4 days in 2% DMEM before transfer to T175 flasks for 6 days prior to harvest and centrifugation of viral stock. Viral titers were determined by serial dilution and quantification of infected foci in Vero E6 cells 24 hpi via immunoperoxidase staining with anti-DENV4 hyperimmune mouse ascitic fluid (HMAF) (ATCC), horseradish peroxidase (HRP)-labeled anti-mouse IgG (1:2,000; KPL-074-1806), and 3-amino-9-ethylcarbazole. ZIKV infections were performed by absorbing virus onto primary hBMEC/hSerC monolayers (2-4 x10^5^ cells) for 2 hours at 37°C, phosphate-buffered saline (PBS) washed, and resupplemented with media. For all experiments involving doxycycline induced, TTP expressing hBMECs and hSerCs, 200 ng/mL doxycycline was maintained in culture media throughout the course of ZIKV infection.

### Induction and Expression Analysis

hBMECs (2 x10^5^) were ZIKV infected, transfected with 0.5 µg/mL Poly (I:C) and Fugene6 (1:3), or treated with 1,000 U/ml IFN⍺ (Sigma) for 24 hours prior to supernatant and RNA collection. For IFN and ISG induction experiments, IFN-⍺ (Sigma) and IFN-λ_1_ generated from IFN-λ_1_ expressing HEK-293T cells, were added to hBMEC/hSerCs concurrently with ZIKV (MOI 0.5). Following inoculum removal, indicated concentrations of IFN-⍺/λ_1_ were added back to monolayers before cell fixation or RNA collection 24 hpi.

### Lentivirus, Transduction, and Selection

pLentiCRISPRv2 plasmid was purchased from Addgene (#52961) and gRNA inserted as described previously (95). pLV[2CRISPR]-hCas9:T2A:Puro-U6 was purchased from VectorBuilder for TTP KO. Lentivirus was produced in HEK 293T cells following PEI transfection (5µg/1µg DNA) of pLentiCRISPRv2, psPAX2 (Addgene #12260), and pLp/VSVG (Invitrogen) and collected 5 days post transfection. hBMECs were doubly transduced over 48 hours and selected with puromycin (0.8µg/mL) for 72 hours prior to KO validation by western blot. gRNAs used can be found in supplementary table 1.

pCW57-MCS1-2A-MCS2 was purchased from Addgene (#71782) and used to conditionally express TTP and IFNλ_1_ derived from ZIKV infected hBMEC cDNA in the presence of doxycycline (0.2–1.0 µg/mL). TTP and IFN-λ_1_ were cloned using primers found in supplementary table 1 and inserted between pCW57-MCS1-2A-MCS2 EcoRI (NEB # R3101S) and BamHI (NEB # R3136S) restriction sites.

### qRT-PCR Analysis

RNA was purified using RNeasy Mini Kit (Qiagen) according to the manufacturer’s instructions. cDNA was generated using Transcriptor First-strand cDNA synthesis kit (Roche) and ProtoScript II first strand cDNA synthesis kit (New England BioLabs) using Oligo-p(dT)_15_ or d(T)_23_ VN primers. qRT-PCR primers used are found in supplementary table 1.

cDNA was amplified using PerfeCTa® SYBR Green SuperMix (QuantaBio) on a Biorad CFX96 Touch Real-Time PCR Detection System. All genes were normalized to internal β-actin controls and gene induction determined using 2^-ΔΔCt^ over uninfected/uninduced controls. For mRNA half-life determination, hBMECs were treated with 5 µg/mL Actinomycin D (Sigma) for the indicated times prior to RNA purification. mRNA abundance over time was plotted relative to RNA collected immediately following actinomycin addition and assessed for significance via one-way ANOVA using GraphPad Prism.

### ELISA

Levels of secreted IFN-β and IFN-λ in mock, ZIKV infected, and Poly (I:C) treated supernatants were determined using DuoSet ELISA (R&D Systems) according to manufacturer’s instructions. ELISA plates (Immunolon 2, U-bottom; Dynatech Laboratories) were developed using tetramethylbenzidine and optical density measured using a Molecular Devices SpectraMax M5 plate reader (450nm). Protein concentrations were calculated by fitting O.D. to a standard curve of purified IFN using SoftMax Pro software.

### Western Blot Analysis

Cells were washed in PBS prior to lysis on ice in buffer containing 1% NP-40, 0.1% SDS, 150 mM NaCl, 50mM Tris-HCl pH 7.4, 2 mM EDTA, 10 nM NaF, 1 mM PMSF, and protease inhibitor cocktail (sigma). Protein concentration was determined using Pierce BCA Protein Assay Kit (Thermo) and 36 µg of protein resolved on a 10-12% SDS polyacrylamide gel. Proteins were transferred to a 0.4µm nitrocellulose membrane and blocked in either 5% bovine serum albumin or 5% non-fat milk prior to incubation with primary antibody. Cellular fractionation was performed as previously described (96). Antibodies used include ZIKV NS5 1:1000 (GeneTex #GTX133327), AUF1 1:1000 (Millipore 07-260), Lamin B1 1:1000 (Cell Signaling D4Q4Z), TTP 1:250 (Cell Signaling D1I3T), IRF3 1:1000 (Cell Signaling D83B9), Myc 1:1000 (Cell Signaling 2272S), KHSRP 1:1000 (Cell Signaling E2E2U), HuR 1:1000 (Santa Cruz sc-5261), GAPDH 1:3000 (Sigma G9545).

### Statistical Analysis

Results in each figure were derived from data collected from ≥3 independent experiments and displayed as the mean ± SEM. One-way analysis of variance, two-way analysis of variance, and unpaired student’s T test were used where indicated, with P-values ≤ 0.05 considered significant. All statistical tests were performed using GraphPad Prism version 9.

**Figure S1 ZIKV induced MXA, IFIT1, and CCL5 are IRF3 dependent** WT hBMECs and hBMECs expressing dominant negative IRF3Δ60 were ZIKV infected (MOI 1) for indicated times prior to qRT-PCR analysis of MXA (A), IFIT1 (B), and CCL5 (C). Data are represented as the mean ± SEM. Experiments were performed 3 times.

**Figure S2: Validation of TTP KO and doxycycline-induced TTP overexpression.** (A, B) WT and TTP KO hBMECs were ZIKV infected (MOI 1). The KO of TTP (A) and overexpression of TTP (B) confirmed by Western blot.

**Table S1: List of primers.** Primers referenced in the methods are detailed.

