## Supplemental Fig 1 for "ZIKV Induction of Tristetraprolin in Endothelial and Sertoli Cells Post-Transcriptionally Inhibits IFNβ/λ Expression and Promotes ZIKV Persistence"

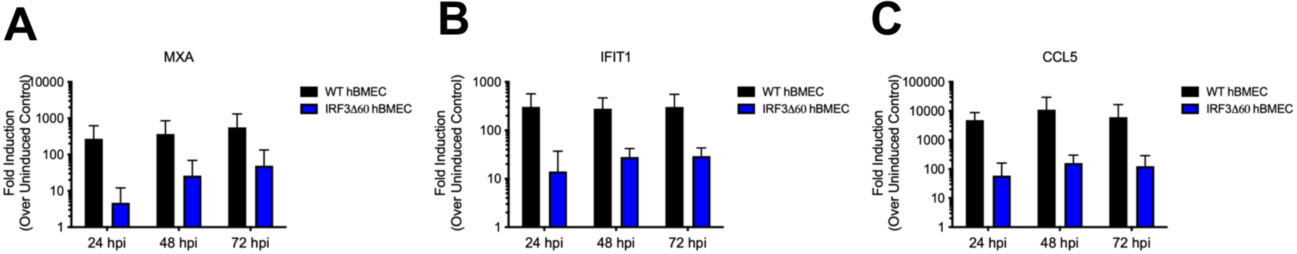
A
MXA
Fold Induction
(Over Uninduced Control)
10000
1000
100
10
1
24 hpi
48 hpi
72 hpi
WT hBMEC
IRF3Δ60 hBMEC
B
IFIT1
Fold Induction
(Over Uninduced Control)
1000
100
10
1
24 hpi
48 hpi
72 hpi
WT hBMEC
IRF3Δ60 hBMEC
C
CCL5
Fold Induction
(Over Uninduced Control)
100000
10000
1000
100
10
1
24 hpi
48 hpi
72 hpi
WT hBMEC
IRF3Δ60 hBMEC

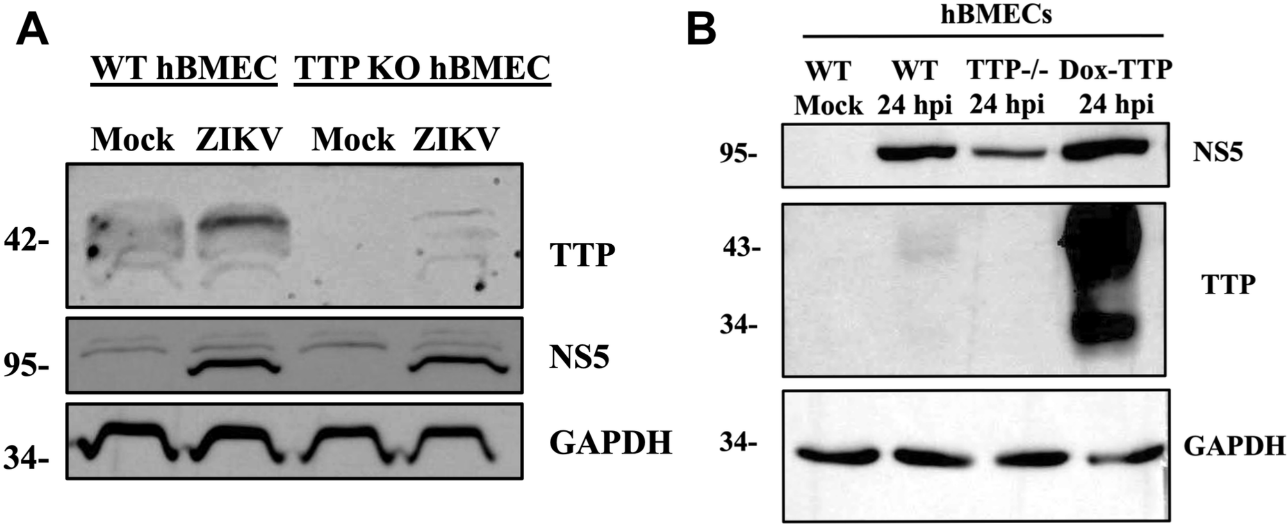

A
WT hBMEC
TTP KO hBMEC
Mock
ZIKV
Mock
ZIKV
42-
TTP
95-
NS5
34-
GAPDH
B
hBMECs
WT
Mock
WT
24 hpi
TTP-/-
24 hpi
Dox-TTP
24 hpi
95-
NS5
43-
34-
TTP
34-
GAPDH

| gRNA | | | |
|---|---|---|---|
| TTP | GTGCCCGTGCCATCCGACCA | AGAGTGGGCGCTGCCGCTAC | AAGTGGCAGCGAGAGCCGTA |
| **Clonging Primers** | | | |
| TTP | Fwd: AAACGTCTCGGAATTCATGGCCAACCGTTACACCATGGATCTGACTG | Rev: AAACGTCTCTGGATCCTCACTCAGAAACAGAGATGCGATTGAAGATGGG | |
| IFN-λ1 | Fwd: AAAGAATTCAGTTGCGATTTAGCCATGGCTGCAGCTTGGACCGTGG | Rev: AAAGGATCCTTCCAATTCCTTGTTGTTTATTTGTGCATAATGTATAAAAGTAAAAC | |
| **qPCR** | **fwd** | **rev** | |
| β-Actin | CACCATTGGCAATGAGCGGTT | AGGTCTTTGCGGATGTCCACGT | |
| IL6 | AGACAGCCACTCACCTCTTCA | TTCTGCCAGTGCCTCTTTGCTG | |
| TTP | CTGTCACCCTCTGCCTTCTC | TCCCAGGGACTGTACAGAGG | |
| IFN-β | AGTAGGCGACACTGTTCGTG | GAGAAGCACAACAGGAGAGCA | |
| IFN-λ1 | CGCCTTGGAAGAGTCACTCA | GAAGCCTCAGGTCCCAATTC | |
| ISG15 | AGGGACACCTGGAATTCGTT | GCGAACTCATCTTTGCCAGT | |
| MXA | TGATCCAGCTGCTGCATCCC | GGCGCACCTTCTCCTCATAC | |
| CCL5 | CGCCTTGGAAGAGTCACTCA | GAAGCCTCAGGTCCCAATTC | |
